# Transgene Expression in Cultured Cells Using Unpurified Recombinant Adeno-Associated Viral Vectors

**DOI:** 10.1101/2023.03.20.533580

**Authors:** Brian Benyamini, Meagan N. Esbin, Oscar Whitney, Nike Walther, Anna C. Maurer

## Abstract

Recombinant adeno-associated viral vectors (rAAV) can achieve potent and durable transgene expression without integration in a broad range of tissue types, making them a popular choice for gene delivery in animal models and in clinical settings. In addition to therapeutic applications, rAAVs are a useful laboratory tool for delivering transgenes tailored to the researcher’s experimental needs and scientific goals in cultured cells. Some examples include exogenous reporter genes, overexpression cassettes, RNA interference, and CRISPR-based tools including those for genome-wide screens. rAAV transductions are less harmful to cells than electroporation or chemical transfection and moreover do not require any special equipment or expensive reagents to produce. Crude lysates or conditioned media containing rAAVs can be added directly to cultured cells without further purification to transduce many cell types – an underappreciated feature of rAAVs. Here, we provide protocols for basic transgene cassette cloning and demonstrate how to produce and apply crude rAAV preparations to cultured cells. As proof-of-principle, we demonstrate transduction of three cell types that have not yet been reported in rAAV applications. We discuss appropriate uses for crude rAAV preparations, the limitations of rAAVs for gene delivery, and considerations for capsid choice. The simplicity of production, exceedingly low cost, and often potent results make crude rAAV a primary choice for researchers to achieve effective DNA delivery.

**Summary:** Recombinant adeno-associated virus (rAAV) is widely used for clinical and preclinical gene delivery. An underappreciated use for rAAVs is the robust transduction of cultured cells without the need for purification. For researchers new to rAAV, we provide a protocol for transgene cassette cloning, crude vector production, and cell culture transduction.

## Introduction

Elucidating the molecular bases of cellular functions often requires the expression of transgenic DNA in cell culture. To be expressed, transgenes must penetrate through a cell’s selective membrane and reach the nucleus. Therefore, the ability to effectively bypass the cell’s physical barriers and manipulate its central processes is a necessity for applying transgenesis to uncover new biological phenomena. One approach capitalizes on the intrinsic ability of viruses to deliver and express foreign DNA.

Adeno-associated virus (AAV) is one of the smallest mammalian viruses: its 4.7kb single-stranded DNA genome contains two genes, *rep* (for replication) and *cap* (for capsid), packaged inside of a 60-mer icosahedral capsid measuring 25nm. The *rep/cap* genes have multiple promoters, reading frames, and splice products which encode at least nine unique proteins required for viral replication, production, and packaging^1,2^.

Additionally, both ends of the genome contain secondary structures called inverted terminal repeats (ITRs) that are necessary for DNA replication, genome packaging, and downstream processing during transduction^3–6^. The ITRs are the only DNA elements that are required for the packaging of the genome into the capsid and therefore AAV can be cloned for transgene delivery purposes by replacing the viral *rep/cap* genes with a researcher’s choice of regulatory elements and/or genes of interest^2^. The resulting recombinant AAV (rAAV), with an engineered vector genome (VG), is widely used in the clinic for human gene therapy and has amassed successes^7^. An underappreciated use of the vector is in the laboratory; rAAVs can efficiently achieve transgene expression in cultured cells to fulfill a researcher’s experimental needs.

The most common method for producing rAAV is by triple-plasmid transfection into HEK293 or 293T cells (Figure 1). The first plasmid, commonly called the *cis* plasmid, contains the desired transgene flanked by ITRs (pAAV). Depending on the application, *cis* plasmids with common elements, such as strong promoters or CRISPR-based tools, are available for purchase. The second is the pRep/Cap plasmid that contains the wildtype AAV *rep* and *cap* genes provided *in-trans*—i.e., on a separate, non-ITR containing plasmid that expresses regulatory and structural elements that then interact with the *cis* plasmid—and is thus called the *trans* plasmid. In addition to physically enclosing the VG, the capsid influences cellular tropism^8,9^. By providing the serotype-specific *cap* gene *in-trans*, researchers are easily able to maximize transduction efficiency by choosing an optimized capsid serotype for their given target cell. Lastly, as a *Dependoparvovirus*, AAV requires a “helper” virus to activate *rep/cap* expression from its viral promoters, achieved by Adenoviral “helper” genes provided on a third plasmid such as pAdΔF6^10,11^. 72 hours after triple-plasmid transfection, the vector can be released from producer cells into the culture media by repeated freeze/thaw cycles. The entire plate contents are then collected, and large cellular debris are removed by centrifugation; the resulting media supernatant is a crude rAAV preparation ready for downstream transductions.

**Figure 1:**
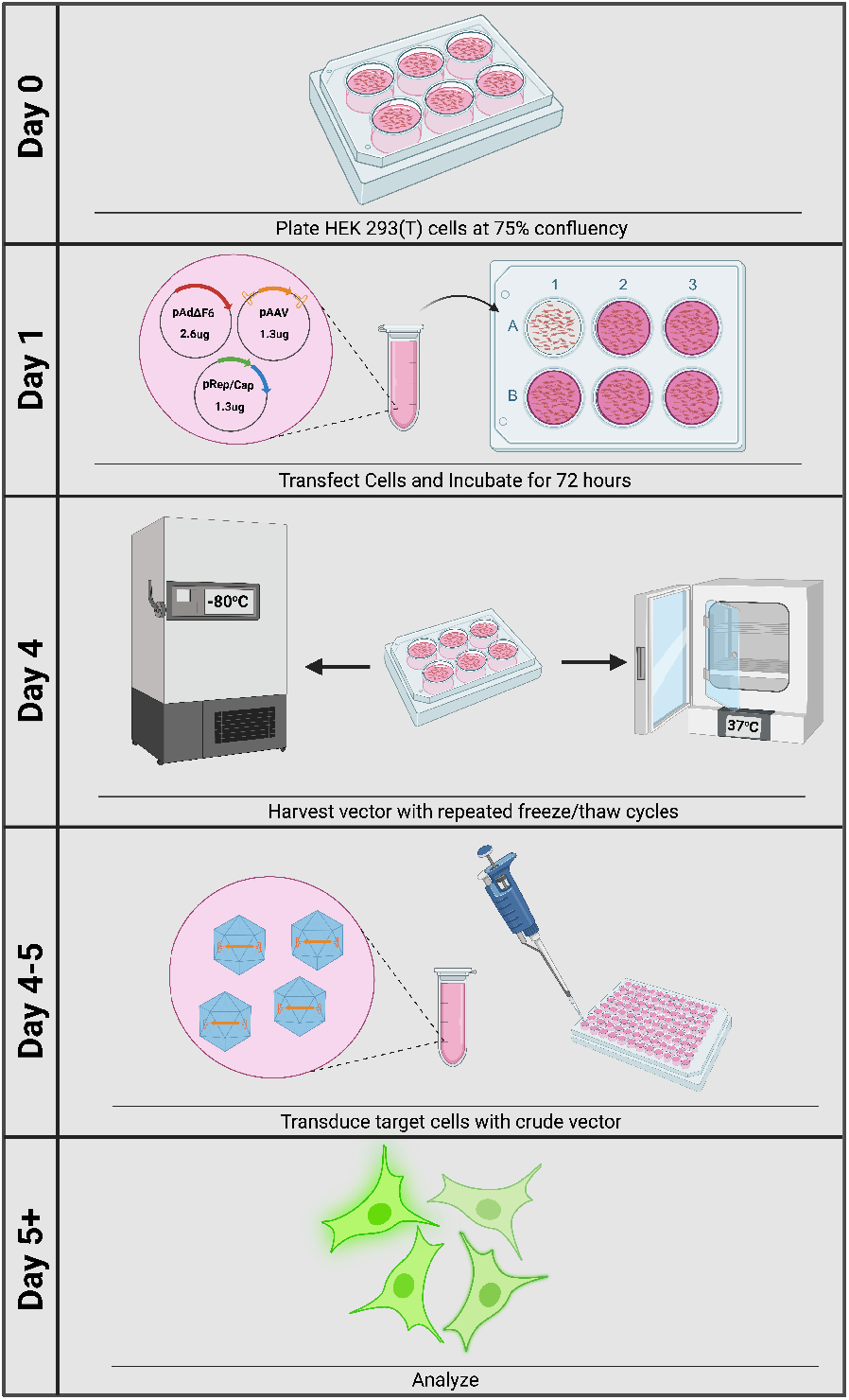
Overview of Crude rAAV Vector Production. Crude rAAV production and transduction can be accomplished within five days.

rAAV may be more favorable for transgene delivery compared to other transfection methods, which are commonly associated with cellular toxicity, low efficiency, and expensive reagents and equipment, such as for electroporation or chemical/lipid-based transfection. rAAV bypasses these obstacles and often provides potent transgene expression with minimal toxicity, and minimal hands-on time. Importantly, producing rAAV and applying it in cell culture is simple and rarely requires purification of the vector from the culture media (Figure 1). Additionally, rAAV does not integrate its VG into the host genome, unlike Lentiviral transgene delivery, and thus lowers the risk of insertional mutagenesis^12^. However, despite the added potential benefits of using rAAV for transgene delivery, limitations must be considered. Importantly, the size of the transgene, including the ITRs, should not exceed 4.9kb due to physical constraints of the capsid, thereby limiting a researcher’s ability to effectively deliver large regulatory elements and transgenes. Furthermore, since rAAV is a non-integrating virus, transduction results in transient transgene expression in dividing cells and may not be practical for stable expression. However, methods using dual rAAV-delivered Cas9 and homology-directed repair (HDR) templates may be used to stably insert sequences at specific genomic loci if a researcher wishes^13^.

## Protocol

### Defined Terms

Virus: an infectious agent that can replicate within an organism
Vector: an agent that transfers genetic material to an organism without replicating
Transgene: a gene that has been artificially added from one species to another
Vector genome (VG): the cloned DNA fragment that contains a transgene flanked by ITRs and delivered by rAAV
Serotype: Natural variations of capsid proteins that are immunologically distinguishable from one another
Capsid Variant: any engineered or synthetic capsid that is distinct in sequence from other capsids but not necessarily formally recognized as an immunologically distinct serotype.
Crude preparation: unpurified vector
Titer: the concentration of vector in a preparation (typically given in units of VG/mL)
Transduction: the process by which transgene expression is achieved in a cell through a vector

#### 1. Required Plasmids

This protocol will clone a gene of interest (GOI) into an ITR-containing plasmid with a CMV promoter and a SV40 poly-adenylation sequence (pAAV.CMV.Luc.IRES.EGFP.SV40, Addgene, Catalog #105533). Plasmids with different regulatory elements or for different applications, such as for CRISPR-based experiments, are available online and will follow similar cloning steps as those below (see discussion for additional cloning steps). Additionally, rep/cap, or *trans*, plasmids for different serotypes can be used—this protocol will use rep/cap for AAV serotype 2.

1.1 Obtain bacterial stabs containing an ITR-containing *cis* plasmid (Catalog #105533), an adenovirus helper plasmid (Catalog #112867), a rep/cap plasmid (Catalog #104963), and a plasmid containing a GOI from Addgene. Streak the bacteria on individual agar plates containing 0.1mg/mL carbenicillin — or other antibiotic depending on plasmid resistance of GOI. Incubate the bacteria at 30° C overnight to grow. Note: if a plasmid with a GOI is not readily available for purchase, a synthetic DNA fragment (gBlock) can be purchased through *Integrated DNA Technologies* (IDT).
1.2 Pick a single colony from each plate and grow in 3mL LB buffer supplemented with 0.1mg/mL carbenicillin at 30° C overnight, shaking at 180 rpm. Transfer 1mL of the bacteria into a sterile 250mL Erlenmeyer flask containing 50mL of LB buffer supplemented with 0.1mg/mL carbenicillin. Shake at 30° C, 180 rpm overnight.
1.3 Transfer the bacteria to a 50mL conical tube and centrifuge for 20 minutes at 3,000 x g, room temperature. Discard supernatant. Isolate the plasmid using a Qiagen Plasmid Plus Midi Kit. Note: ITRs are unstable structures—to prevent ITR deletions or mutations, bacterial cells should be grown at 30° C to slow cellular division and mitigate errors in replication. To increase plasmid yield and purity, a midiprep or maxiprep kit is ideal. It is recommended to use an endotoxin-low or -free kit (such as Qiagen Plasmid Plus Midi Kit) to reduce downstream endotoxin contamination of cells.

#### 2. Cloning Gene of Interest into AAV ITR-containing Plasmid

2.1 Open the plasmid map containing the GOI in a DNA viewing software, such as Snapgene. Note: The full cassette, including the ITRs, should not exceed 4.9kb due to physical packaging limitations of the protein capsid. Additionally, PCR amplification cannot be achieved through ITRs due to secondary structures. As such, Gibson assembly is not recommended for rAAV transgene cloning.

2.1.1 Create a forward primer that extends from the 5’ most region of the GOI inward to the gene body, until a Tm of ~55 °C has been reached. Include the sequence for a EcoRI restriction site (5’ GAATTC 3’) at the 5’ most end of the primer sequence. Additionally, add six extra bases 5’ of this restriction site to allow for the enzyme to efficiently interact with the DNA.
2.1.2 Use an online primer design tool, such as *Oligo Calc*, to ensure no hairpins or self-complementary sequences are present in the primer.
2.1.3 Repeat the steps to create a reverse primer at the 3’ most region of the GOI inward to the gene body, while instead including a NotI restriction site (5’ GCGGCCGC 3’). Note: Your GOI cannot contain EcoRI or NotI restriction sites—if so, different restriction enzymes will need to be used.
2.2 Amplify the GOI in a 50uL PCR reaction. Use the required components from Table 1 (adapted from NEB).

2.2.1 Place the reaction in a thermocycler and follow the parameters from Table 2 for programming. If primers have different Tm, use the lower Tm for the program.
2.2.2 Verify amplification of correct PCR fragment by adding 1uL gel loading dye (6x) to 5uL PCR reaction. Run a DNA ladder and this mixture on a 0.8% agarose gel containing ethidium bromide and visualize with UV light. Caution: Ethidium Bromide is a known mutagen.
2.2.3 If a single, specific band appears on the gel, purify the remaining PCR product in the PCR strip tube using a DNA purification kit. If multiple bands appear (due to non-specific amplification), excise the desired band, and purify the DNA using a Gel Extraction Kit.
2.3 Digest the purified PCR product and pAAV.CMV.Luc.IRES.EGFP.SV40 backbone plasmid in a 50uL reaction with the required components from Table 3 for 1hr at 37 °C. A larger quantity of pAAV plasmid DNA is required due to expected inefficient recovery following gel extraction. Note: The restriction enzymes used are High Fidelity (HF) versions. Regular NotI cannot be used with CutSmart buffer. If different enzymes are being used, a different incubation temperature and buffer may be required.

2.3.1 Purify the digested PCR product with a DNA purification kit. Prepare the digested pAAV.CMV.Luc.IRES.EGFP.SV40 backbone plasmid for gel electrophoresis by adding 10uL gel loading dye (6x) to the 50uL digest reaction. Load a DNA ladder, digestion reaction, and 250ng undigested plasmid (negative control) on a 0.8% agarose gel containing ethidium bromide with wide-teeth wells. Visualize with UV light, excise the desired fragment (~4.5 kb), and purify the DNA using a Gel Extraction Kit.
2.4 Ligate the digested PCR product and pAAV backbone plasmid in a 20uL ligation with the required components from Table 4. To calculate the amount of PCR product needed, use Formula 1. Typically, a 3:1 ratio of PCR product (insert) to backbone (pAAV), with a mass of 50ng backbone, is used.

2.4.1 Perform a negative control by replacing the PCR product with water. Incubate ligation reactions at 16 °C overnight, or at room temperature for 2 hours.

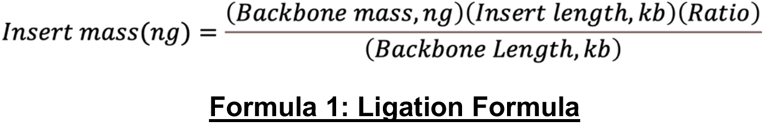
2.5 Thaw a vial of competent, recombination-deficient bacterial cells (such as Stbl3 cells) on ice and place 50uL into a 1.5mL Eppendorf tube. Add 3uL of the ligation reaction directly onto the cells and incubate on ice for 30 minutes. Note: ITRs are unstable structures and therefore require recombination-deficient bacterial strains to limit ITR deletion and recombination events. DH5α cells may be used intermittently for propagation (one or two times) but are not recommended for long-term use. ITR integrity should be checked regularly after midiprep purification.

2.5.1 Heat-shock the bacteria in a 42° C water bath for 30 seconds then immediately transfer to ice for 2 minutes. Add 200uL LB buffer and shake for 1 hour at 37° C, 180 rpm.
2.5.2 Plate 125uL of the mixture onto an agar plate containing 0.1 mg/mL carbenicillin and incubate overnight (~18 hours) at 30° C. Pick multiple clones and place each into a sterile plastic culture tube with 3mL LB containing 0.1mg/mL carbenicillin. Incubate overnight shaking at 30° C, 180 rpm. Note: ITRs are unstable structures—to prevent ITR deletions or mutations, bacterial cells should be grown at 30° C to slow cellular division and mitigate errors in replication. Additionally, smaller colonies should be picked as those without ITRs may have a growth advantage over others.
2.5.3 Pipette 1.8mL of each culture into a 2mL tube and centrifuge at 6,000 x g for 3 minutes. Isolate the DNA using a miniprep kit. Store the remaining 1.2mL at 4° C. Verify correct clones by DNA sequencing or diagnostic restriction enzyme digest. Note: Sequencing is inefficient through ITRs due to secondary structures; if desired, use primers that will not sequence through the ITRs
2.5.4 Add 50mL LB buffer containing 0.1mg/mL carbenicillin to a sterile 250mL Erlenmeyer flask. Add 500uL of the remaining culture to the flask and shake at 30° C, 180 rpm until an OD600 of 2-3 is reached (~18 hours). Transfer the bacteria to a 50mL conical tube and centrifuge for 20 minutes at 3,000 x g, room temperature. Isolate the DNA using an endotoxin-low or -free midiprep kit.
2.5.5 Check ITR integrity with a 20uL diagnostic restriction enzyme digest using *XmaI* (or *SmaI*). The reaction should contain 500ng plasmid, 0.5uL enzyme, and 1x buffer; incubate for 1 hour at 37° C. Run the reaction on a 0.8% agarose-gel. If ITRs are intact, digestion will give a band at ~2.9kb and another band the size of your cassette. If this distinct band is not observed, the ITR may not be intact and cloning will need to be redone. Note: Plasmids containing *XmaI* (or *SmaI*) restriction sites (in addition to those in the ITRs) will have additional bands appear on the gel.

**Table 1:** PCR amplification reagents.

| Amount | Component |
| --- | --- |
| X uL | Template DNA (25 ng) |
| 1 uL | F primer (10 uM) |
| 1 uL | R primer (10 uM) |
| 1 uL | dNTP (10 mM) |
| 10 uL | Phusion Buffer (5x) |
| 1 uL | Phusion Polymerase |
| X uL | Water |
| 50 uL | Final Volume |

**Table 2:** PCR amplification programming.

| Step | Temperature | Time |
| --- | --- | --- |
| Initial Denaturation | 98° C | 1 minute |
| 30 Cycles | 98° C | 10 seconds |
|  | T<sub>m</sub> of Primer | 30 seconds |
|  | 72° C | 30 seconds per kb |
| Final Extension | 72° C | 10 minutes |
| Hold | 4° C | indefinitely |

**Table 3:** Digestion reagents.

| Amount | Component |
| --- | --- |
| X uL | PCR product (1ug) or pAAV plasmid (3ug) |
| 5 uL | CutSmart Buffer (10x) |
| 1 uL | EcoRI-HF |
| 1 uL | NotI-HF |
| X uL | Water |
| 50 uL | Final Volume |

**Table 4:** Ligation reagents.

| Amount | Component |
| --- | --- |
| X uL | Insert (X ng) |
| X uL | Backbone (50 ng) |
| 2 uL | T4 DNA Ligase Buffer (10x) |
| 1 uL | T4 DNA Ligase |
| X uL | Water |
| 20 uL | Final Volume |

#### 3. Vector Production with Triple-Plasmid Transfection

The following values are optimized for a single well of a 6-well plate that yields a final crude preparation volume of 2mL. All values can be scaled up by 10x for a 15cm plate that yields a final volume of 20mL or scaled down by 4x for a 24-well plate with a final volume of 500uL.

3.1 Seed HEK293 or HEK293T cells into a 6-well plate with pre-warmed DMEM (4.5g/L glucose, 110mg/L sodium pyruvate) supplemented with 10% FBS. Grow to ~75%-90% confluency in an incubator set at 37° C and 5% CO2 (Figure 2). Note: Cells cultured with Penicillin/Streptomycin (P/S) have been observed to decrease the vector production yield. It is recommended to remove P/S from cultured cells a few passages (~2) prior to transfection.
3.2 In a 2mL tube, prepare a mixture containing 1.3 ug pAAV2/2 (Rep/Cap, serotype 2), 1.3 ug pAAV.GOI, 2.6 ug pAdΔF6, and serum-free (SF) DMEM (4.5g/L glucose, 110mg/L sodium pyruvate) into a total volume of 100uL (see excel sheet for convenient calculations). Prepare a negative control in a separate tube by replacing pRep/Cap with any unrelated plasmid. The negative control accounts for any residual plasmid present in the crude preparation that could possibly transfect cells during transduction, although rare. Note: Endotoxin-low or -free maxiprep or midiprep plasmid is ideal for triple-transfection and generally produces higher titer vector compared to miniprep DNA, although miniprep DNA can be used for quick-and-dirty preliminary testing.
3.3 Make a 1ug/uL stock of PEI (polyethylenimine hydrochloride) MAX by dissolving 100mg PEI MAX in 100mL distilled water. Adjust pH to 7.1 with NaOH. Filter sterilize the mixture with a 0.22um filter and freeze 1mL aliquots at −80° C for long-term storage. Once thawed, the reagent can be stored for one month at 4° C.
3.4 Add 5.2uL of the PEI Max to the plasmid mixture—this is a Plasmid:PEI ratio of 1:1. Immediately mix well but gently by pulsing 10-15 times on a vortex mixer set to 7. If multiple vector preparations are produced, add PEI to each plasmid mixture at staggered time intervals of one minute (i.e.—preparation 1 at t = 0 min, preparation 2 at t = 1 min, etc…) to allow for sufficient timing to aspirate wells and dilute the reaction in the following steps. Note: A 1:1 ratio of Plasmid:PEI is the optimal ratio in our hands. Individual users should optimize this ratio for maximum efficiency. Refer to the discussion for how to perform PEI optimization.
3.5 Incubate each tube for exactly 15 minutes and then dilute the reaction with 1.9mL of SF DMEM (4.5g/L glucose, 110mg/L sodium pyruvate) for a final volume of 2mL. <u>Gently</u> pipette twice to mix. Overmixing will disrupt Plasmid:PEI complexes and result in lowered transfection efficiency. 3.6 Aspirate media from well and add Plasmid:PEI mixture gently on well sides to prevent cellular detachment. Incubate cells for 72 hours at 37° C and 5% CO2. Note: Cell lysis and detachment during incubation is a normal process of rAAV production (Figure 3).

**Figure 2:**
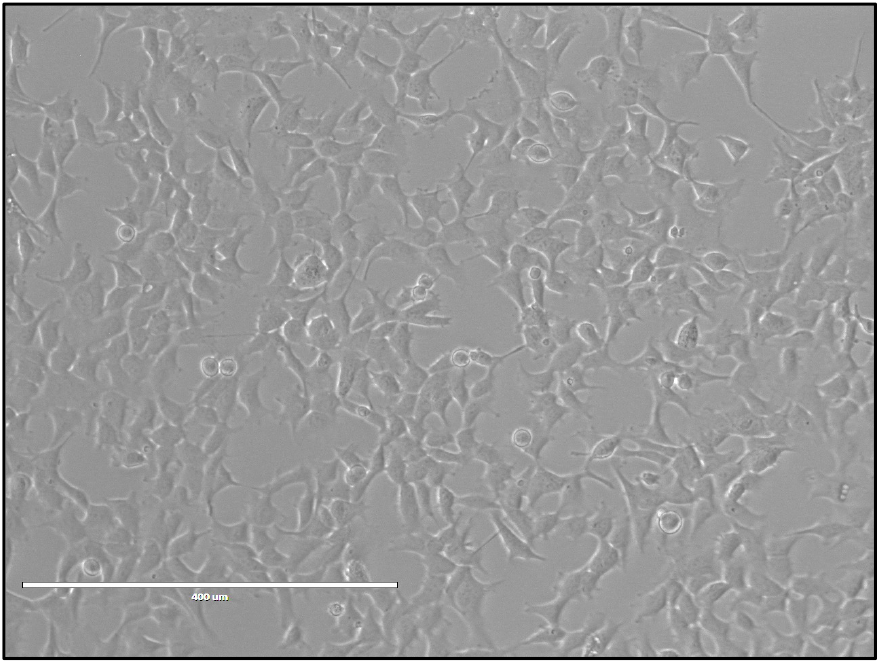
HEK293 cells ready for transfection. The ideal confluency (75-90%) of HEK293 cells needed for triple-plasmid transfection

**Figure 3:**
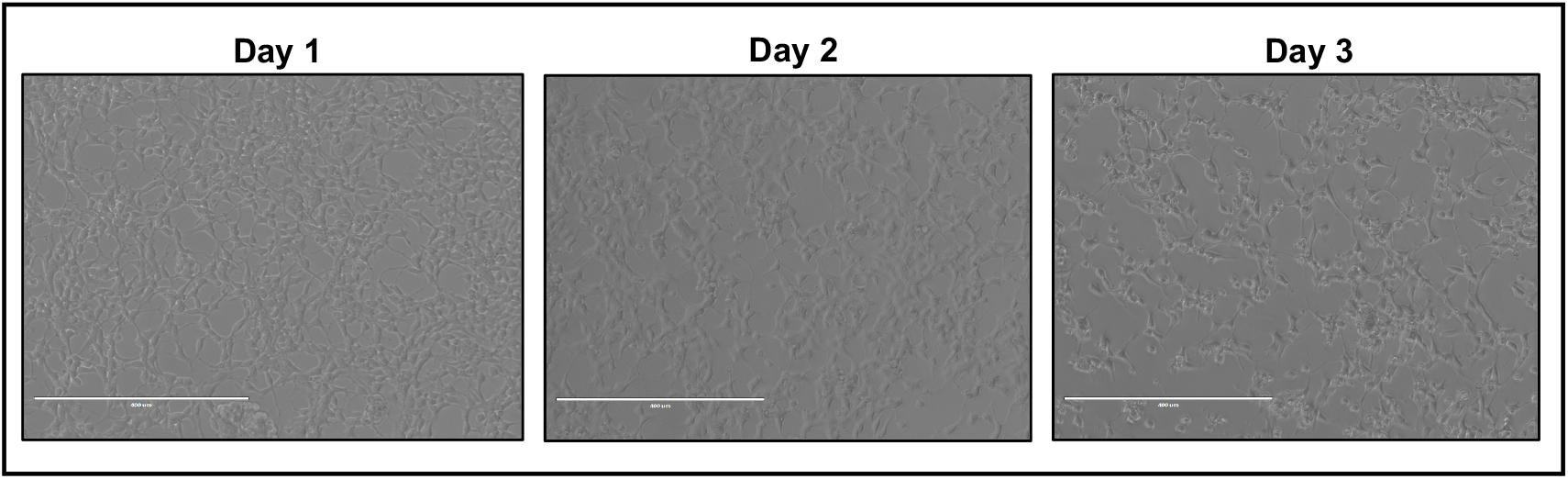
HEK293 cells after transfection. The appearance of HEK293 cells after one, two-, or three-days post transfected with pAAV.CMV.Luc.IRES.EGFP.SV40, pAAV2/2, pAdΔF6, and 1:1 ratio of Plasmid:PEI MAX. Scale bars: 400um

#### 4. Harvesting Crude Vector Preparations

4.1 Freeze the entire plate of transfected cells for 30 min at −80° C, followed by a thaw for 30 min at 37° C. Repeat for a total of 3 freeze/thaw cycles. Do not remove the lid – plate must remain sterile inside. Note: A non-humidified incubator works best for thawing, which minimizes condensation on the outside of plates that can lead to accidental contamination. Plates can remain at −80° C until ready to proceed to next steps.
4.2 Mix each well by pipetting to ensure maximum disruption of cells. Transfer lysate to a 2mL tube and centrifuge for 15 min at 15,000 x g, room temperature to remove cell debris. Note: Steps 2 and 3 should be performed in a laminar flow hood using aseptic techniques.
4.3 Carefully transfer supernatant to a new 2mL tube. This crude preparation can be used immediately without titration or purification. Vector can be stored at 4° C for several months or −20° C for years. Note: If necessary, titers may be calculated by qPCR by quantifying the number of vector genomes (VG) inside of Dnase-resistant particles. As such, titers are given in units of VG/mL. Vector stored for several months at 4°C should be re-titrated before use as vector can aggregate and/or bind tube walls, reducing the titer of the preparation.

#### 5. Transduction

5.1 Plate your desired cell type in a 96-well plate at a target confluency of 50-75%, depending on the duration of transduction. Greater confluency may result in reduced AAV transduction efficiency.
5.2 Vector can be added to cells without knowing the titer of the crude preparation for applications where dose is not relevant, such as testing a panel of capsids for transduction in a new cell type. Perform a 1:3 dilution series of the crude preparation in SF media to obtain the optimal amount of vector required for your application. Aspirate media from 96-well plate and add 100uL diluted crude preparation to wells. Incubate cells at desired temperature. Note: Serum may contain antibodies that can neutralize the AAV vector and reduce transduction efficiency. Serum may be used during transduction for sensitive cell types.
5.3 Vector can be removed and replaced with fresh serum-containing media as soon as 2 hours posttransduction (hpt). For many applications, overnight incubation is more than sufficient to transduce cells and wells can be replaced with fresh media in the morning. Robust cell types, such as U2-OS, can be reliably assayed without any media change.
5.4 Terminate transduction based on your application (e.g.—lysis, fixation, etc.). For most cell types, peak expression occurs by 48hpt and is thus a common experimental endpoint.

## Discussion

### Cloning

The cloning protocol is not limited to the pAAV.CMV.Luc.IRES.EGFP.SV40 plasmid used above and can be easily altered based on a researcher’s experimental needs. Many ITR-containing plasmids are readily available online for purchase. For example, plasmids containing both Cas9 and an sgRNA cloning site are available (Addgene, Catalog #61591), but require few additional steps such as oligonucleotide annealing and PNK treatment. Additionally, plasmids containing a multiple cloning site (MCS) with only ITRs and no inner regulatory elements can be found (Addgene, Catalog #46954). If different plasmids are to be used, the restriction enzymes (RE) used for digestion are typically the only elements that may need to be changed in this protocol.

When isolating plasmid from bacteria, it is recommended to use an endotoxin-low or -free midiprep or maxiprep kit to mitigate harm to cells during triple-plasmid transfection or transduction. Plasmid from miniprep kits often contain higher impurities, reduced concentrations, and fewer supercoiled DNA, all of which can affect the downstream production of rAAV and is thus not recommended.

The structure and properties of ITRs must be considered during cloning. First, it is extremely difficult to PCR through the ITR. Cloning designs that require PCR amplification through ITRs should be avoided, and additionally limits the use of the Gibson Assembly cloning technique. As such, restriction enzyme cloning is the preferred method for cloning into ITR-containing plasmids. Furthermore, certain primers for sanger sequencing may not be compatible if the sequenced region contains the ITR. Instead, it is recommended to use primers that sequence away from the ITRs and into the vector genome body to get more precise sequencing results. Second, ITRs are prone to deletions, rearrangements, and mutations when transformed into bacteria for plasmid amplification^14,15^. To mitigate these events, it is recommended to use recombination-deficient competent bacterial strains, such as Stbl3, and to incubate them at 30° C to slow down cellular divisions. Last, we have observed that smaller colonies may correspond with clones without rearrangements or deletions, as those without ITRs may confer a growth advantage and be larger. Therefore, it is recommended to pick colonies that are small.

### Vector Production

The successful production of rAAV vector can be affected by multiple elements. One important factor is the health of HEK293 or 293T cells used for transfection. Generally, low passage numbers are ideal, as highly passaged cells may exhibit genotypic and phenotypic variances that can reduce rAAV titers. Additionally, the density of the seeded cells should be 75-90% confluency for effective production. Sparse cells generate low vector yields because there are less cells available to produce vector, while overgrown cells will not be efficiently transfected.

Variations between reagent lots, cell stocks, and general lab-to-lab variability contribute to differences in transfection efficiencies and production titer. One optimizable factor that can lead to titer improvements is the Plasmid:PEI ratio in transfection reactions. We recommend a Plasmid:PEI ratio of 1:1 as a starting point, and if transfection or transduction efficiency appears poor, we recommend testing several different ratios. Titer optimization is easiest if using a transgene with a visual readout, such as a fluorescent protein that can later be assayed after transduction. To perform the optimization, follow the steps from “Vector Production with Triple-Plasmid Transfection”, while instead using a 12-well plate and scaling down the plasmid masses and reagent volumes by two (final plasmid mass is 2.6ug). Adjust the PEI volume accordingly to correspond to ratios ranging from 1:0.75 to 1:3, with increasing increments of 0.25 (Figure 4). Dilute each reaction with 950uL SF media after 15 minutes. For convenience, a master mix containing the triple plasmids can be made and individually pipetted into 1.5mL tubes prior to adding PEI—see attached excel sheet titled “PEI optimization for 12-well”. Harvest the vector, transduce cells of interest, and image. The well with the highest transduction efficiency (proportion of GFP+ cells) corresponds to the highest titer and most optimal ratio of PEI:DNA.

**Figure 4:**
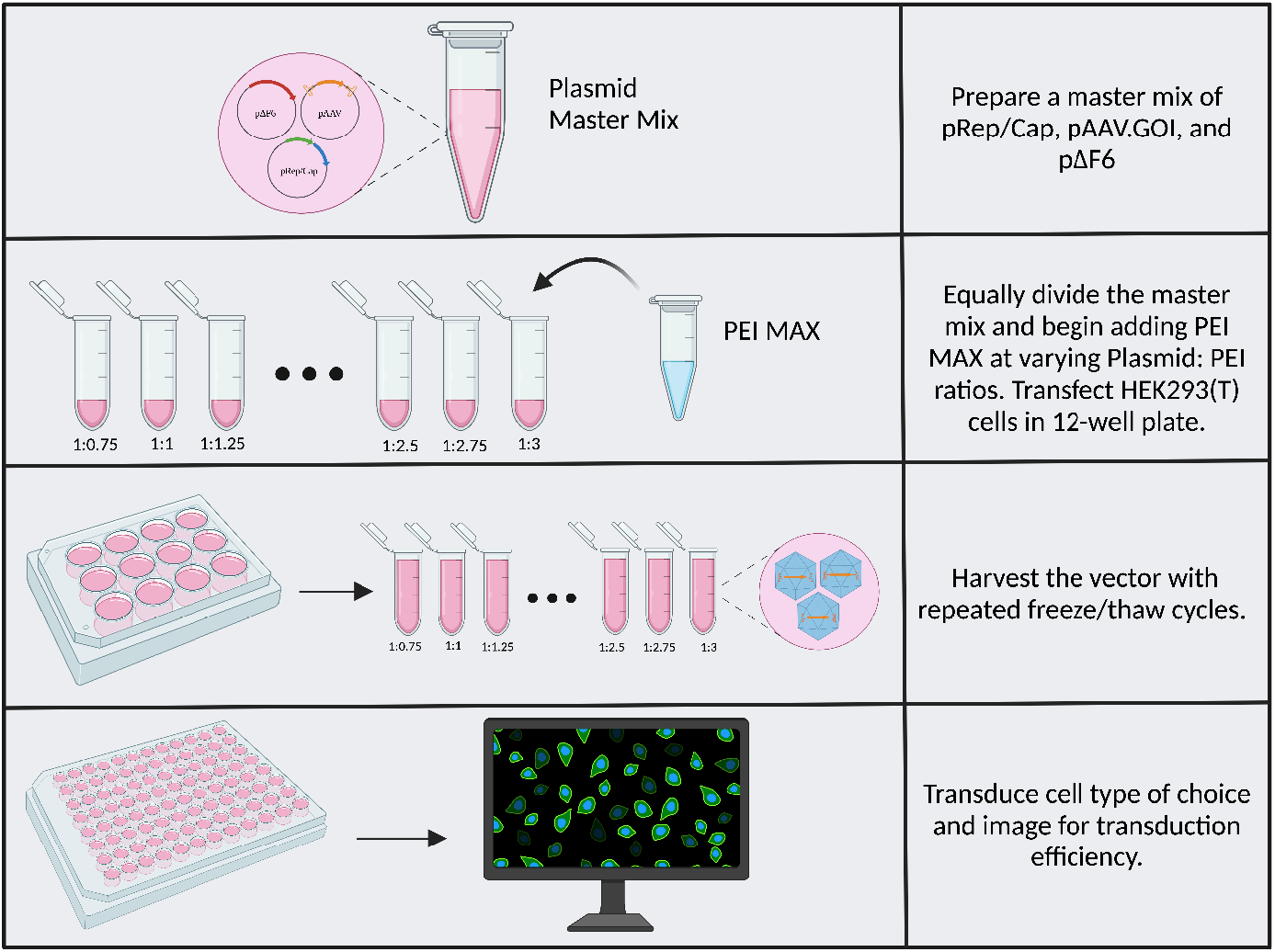
PEI Optimization Workflow.

### Harvest and Titer Considerations

The freeze/thaw technique used to harvest rAAV vector effectively lyses HEK293 cells in a manner compatible with the direct use of the clarified lysate to transduce cultured cells. Certain rAAV serotypes, such as AAV1, AAV8, and AAV9 are released from cells during vector production and can be harvested from the cultured cell medium without freeze/thaw cycles^16^. The method described herein typically yields titers on the order of 1×10^10^VG/mL when using AAV2 capsids, and 1×10^11^ VG/mL for AAV8. While higher titers can be achieved by detergent or other chemical-based lysis, these are harmful to cells in downstream use and require rAAVs to be further purified from the lysate. Lower titer is one tradeoff a researcher should consider when determining whether crude preparations are appropriate for their research needs, however, the marginally lower titers produced by the methods described here can transduce many cell types very well (see representative results). In addition to transfection efficiency and cell health, vector titers vary depending on the capsid serotype used during rAAV production and the size and sequence of the transgene within the VG^17^.

When harvesting crude vector preparations, plasmid DNA that was used during triple-plasmid transfection may be present and, although rare, result in downstream transfection during transduction. Furthermore, unpackaged VGs may bind to the exterior of capsids and invoke an innate immune response to naked and foreign single-stranded DNA^18,19^. Therefore, sensitive cell types may require vector preparations to be DNase digested and purified to remove unpackaged VGs and plasmid.

If one wishes to calculate the titer of a crude preparation, quantitative PCR (qPCR) can be performed to quantify the number of packaged VG inside DNase-resistant particles (DRP). Briefly, a small amount of crude preparation is Dnase-digested to remove plasmid DNA, contaminating nucleic acids, or partially packaged VG. The sample is then subject to qPCR and the protected VG inside of DRPs is quantified, resulting in a titer with units of vector genome per mL of crude preparation^20^. It is not recommended to perform vector titration using ELISA-based assays that quantify <u>capsid</u> titers. Compared to wild-type AAV virus, rAAV suffers from a proportion of empty and partially packaged capsids^21^. ELISA will quantify all capsids regardless of their genome contents and will overestimate the transducible units present in a preparation, which requires a packaged VG.

### Transduction Considerations

Many factors influence rAAV transductions and proper considerations should be made for any new experiment. Depending on the promoter driving transgene expression, expression onset can occur as early as 4 hours post-transduction (hpt), and peak expression is typically achieved by 48hpt. It is important to keep in mind the duration of time from the initial seeding of cells to the experimental endpoint. This is to estimate the starting confluency of the cells and ensure that they do not overgrow by the end of the experiment. If cells become overconfluent, cellular behavior may be altered due to a stress response and can confound experimental results. Some cell types, like U2-OS, can tolerate overgrowth/contact inhibition quite well. Additionally, they can withstand long periods (48hrs+) in serum-free “conditioned” medium – the product of this production protocol. However, sensitive cell types may require serum addition or dilution of the crude preparation with special growth medium to maintain health during transduction. A slightly reduced transduction efficiency from using serum-containing media is a potential tradeoff for cell health and should be considered by the researcher.

Typically, for rapidly dividing cells, a starting confluency of around 50% is optimal for applications that will be terminated 48 hpt. However, confluency can be adjusted accordingly based on the needs of the experiment. It is not recommended to transduce monolayer-type immortalized cell lines over 75% confluency due to decreased transduction efficiencies. Most cultured cell types are successfully transduced and healthy after overnight incubation with crude rAAV preparations, followed by a change to fresh serum-containing media in the morning.

Capsid serotype is an important factor to consider when producing rAAV to transduce a target cell, as the capsid is the primary determinant of cellular tropism and subsequent transgene expression^8^. AAV2 is a widely used serotype due to its ability to effectively transduce many types of cultured cells^9^. This property of AAV2 may be attributed to heparin sulfate proteoglycans (HSPGs) serving as the primary attachment factor for AAV2 and the high levels of HSPGs on cultured cells from the adaptation to growing in a dish^22^. Other capsids, such as AAV9, are less effective at transducing broad cell types and may be explained by their reliance attachment factors that are not expressed in this setting^23^. Therefore, we recommend AAV2 as a first-choice capsid in cultured cells if a desired target cell has not been previously tested with rAAV in the literature.

### Transgene Expression and Potential Integration Considerations

rAAVs do not reliably result in permanent expression of the transgene. Overtime, VGs can become silenced and transgenic expression may be shut down following several passages^24^. Additionally, the majority of VGs remain episomal, and rAAVs do not contain the viral *rep/cap* proteins that would (1) mediate frequent integration into the host genome as in a wild-type viral lysogenic infection or (2) promote replication of VGs^25^. As a result, episomes in transduced cells will eventually be diluted out among daughter cells through divisions.

Basal-level integration is a possibility for all delivered transgenic DNA material. However, ITR-containing VGs are prone to integration at a higher frequency^26^. Therefore, permanent expression of a transgene may be observed in a small subset of cells. Users should consider this possibility especially when using rAAV to deliver DNA-cutting enzymes, such as Cas9, as double-stranded breaks may result in an even larger frequency of integration and permanent expression^27^. While this makes rAAV a good candidate for delivering HDR templates for endogenous tagging or gene addition, the possibility of Cas9 insertion should be considered^13,28^.

## Representative Results

### Finding an optimal capsid for transducing cultured cells of interest

A variety of cell types were transduced to determine the tropism of various capsid serotypes (Figure 5A). The natural serotypes AAV2, AAV4, AAV5, AAV8, AAV9, and the engineered capsid variants Anc80, LK03, DJ, and KP1 were packaged with a vector genome expressing mScarlet under a CMV promoter, cloned using the methods provided in this protocol. Untitrated crude vector preparations were diluted in the appropriate culture media and cells were transduced for 24+ hours and imaged (Figure 5B). Transduction efficiency was calculated by the proportion of total cells that were mScarlet+, or the proportion of cells in a single *z*-plane that were mScarlet+ (for mouse small intestinal organoids).

**Figure 5:**
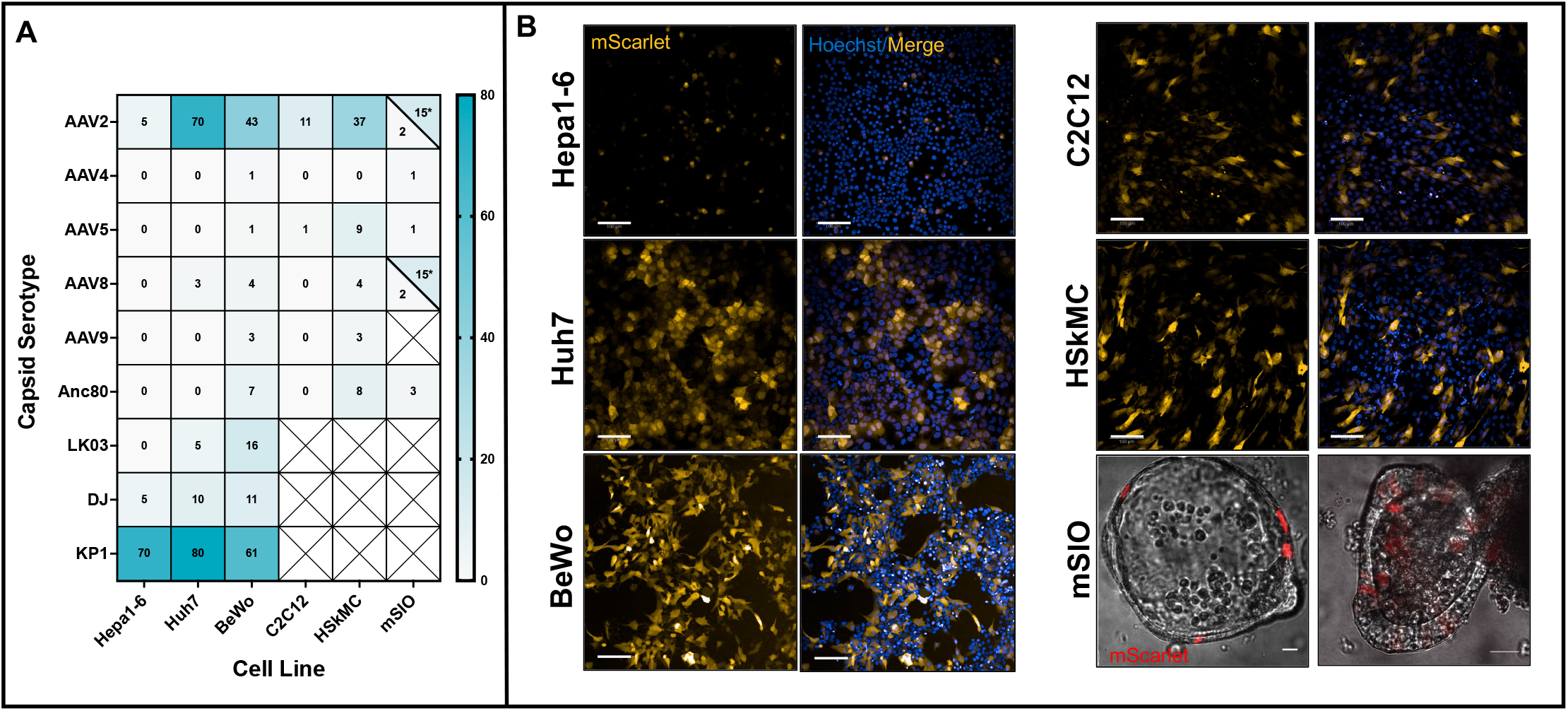
Crude Vector Transduction of Various Cell Types. (A) Cells were treated for 24+ hours with undiluted (Hepa1-6 and Huh7) or 50% diluted (BeWo, C2C12, HSkMC, and mSIO) crude preparation containing a CB7.mScarlet expression cassette (~2.6kb) packaged inside the associated AAV capsid serotype. Mouse small intestinal organoids (mSIOs) were either AAV treated in pre-transduction media or organoid growth media*. Values are given as percent of total cells that are mScarlet+. (B) AAV2 crude vector preparations packaging CB7.mScarlet were added to cells for 24+ hours and imaged using mScarlet and Hoechst filter sets. mSIOs were either cultured in pre-transduction medium (left) or organoid growth medium (right). Scale bars: 100um (Hepa1-6, Huh7, C2C12, and HSkMC), 200um (BeWo), 20um (mSIO)

AAV2 and KP1 were the most potent serotypes across cell lines tested (Figure 5A). We observed that similar cell types originating from different species are transduced at varying efficiencies. For example, Hepa1-6 (a murine-derived liver cell line) exhibits a ~15-times less effective transduction compared to Huh7 (a human-derived liver cell line) when using AAV2. Moreover, different media conditions play an important role during transduction. Mouse small intestinal organoids (mSIOs) cultured in pre-transduction media are less effectively transduced compared to those cultured in organoid growth media.

These results show that crude vector preparations can be used effectively to transduce a variety of cell types without further steps such as purification and titration. However, careful consideration should be taken when choosing a capsid to maximize transgene delivery. Many other cell lines and capsids have been previously tested and are published, though with purified vector preparations^9^. Nonetheless, crude vector preparations generally exhibit the same trends in efficiency and can be used in a similar manner.

### Mouse small intestinal Organoids

Mouse small intestinal organoids (mSIOs) were derived from a small intestinal crypt prep from C57BL/6J mice. Organoids were embedded in 90% Matrigel and cultured in 24-well plates containing either organoid growth medium (without antibiotics) or pre-transduction medium (50% Wnt3a-conditioned medium (produced in-house using L Wnt-3A cells in organoid growth medium), 10 mM nicotinamide, 10uM ROCK inhibitor, and 2.5uM CHIR99021) for one passage (5-7 days) before transduction at 37°C and 5% CO2. Prior to transduction, organoids were rinsed with D-PBS, Matrigel domes were disrupted by pipetting, and organoids were dissociated into small cell clusters by incubating them in TrypLE Express for 10 min at 37 °C and further pipetting. The cell dissociation process was stopped by adding 5% FBS in DMEM/F-12 with 15 mM HEPES, cell clusters were transferred into centrifuge tubes and cell clusters were collected by centrifugation for 5 min at 1000xg. Pelleted cell clusters were resuspended in transduction medium (pre-transduction medium containing 10 μg/mL polybrene or organoid growth medium containing 10 μg/mL polybrene) and transferred into 48-well plates, followed by the addition of AAV crude preps for transduction. Transduction of organoids was performed by centrifuging the plate for 1 h at 600xg and 37 °C and then incubating at 37 °C for an additional 6 h. Transduced organoids were then collected in 1.5 mL microcentrifuge tubes, centrifuged for 5 min at 1000xg, and put on ice. After removal of the supernatant, transduced organoids were resuspended in 90% Matrigel in pre-transduction medium or organoid growth medium and droplets of 20uL were seeded into one well of a prewarmed chambered cover glass. After incubation for 10 min in the incubator, the droplet was embedded in 350 μL of pre-transduction medium or organoid growth medium and incubated at 37°C in the incubator. The transduction efficiency was determined 2-5 days after transduction by imaging the transduced organoids on a Zeiss LSM900 confocal microscope (Plan-Apochromat 40x/1.3 Oil DIC (UV) VIS-IR M27 objective; excitation with a 561 nm laser; detection of red fluorescence with a GaAsP-PMT detector and brightfield detection with a photodiode; bidirectional line scanning; acquisition of representative single *z* planes) and estimating the percentage of red fluorescent cells per organoid.

### BeWo cells

Human placental choriocarcinoma BeWo cells were plated at 10,000 cells per well of 96-well plate 24hrs before transduction. Crude AAV preps were then diluted 1:1 in serum-free F12-K BeWo medium up to a total volume of 50uL. Cells were incubated at 37°C overnight (~18hrs) and then 100uL F12-K BeWo medium containing 10% FBS was added to bring up the total volume to 150uL. Cells were incubated an additional 6hrs for a total of 24hrs post-transduction. For imaging, live cells were stained with Hoechst dye for 30min in F12-K medium, then the medium was exchanged for phenol-free DMEM containing 10% FBS, 1x Glutamax supplement, and 1x Sodium Pyruvate supplement for imaging. Imaging was conducted on the Perkin Elmer Opera Phenix using the DAPI (for Hoechst imaging) and mCherry (for mScarlet imaging) filter sets with 37C and 5% CO2.

### Hepa1-6 and Huh7 cells

Murine Hepa1-6 liver cells and Human Huh7 liver cells were plated in DMEM (4.5g/L glucose, 110mg/L sodium pyruvate, 10% FBS) at 5,000 cells per well of a 96-well plate 24hrs before transduction. 50uL of undiluted crude AAV preps were then added directly to cells and incubated at 37°C for 48 hours. Cells were washed once in PBS, fixed in 4% PFA, and stained with Hoechst dye for 10min. Imaging was conducted on the Perkin Elmer Opera Phenix using the DAPI (for Hoechst imaging) and mCherry (for mScarlet imaging) filter sets.

### C2C12 and HSkMC cells

Murine C2C12 myoblasts and Human Primary Skeletal Muscle Cells (HSkMC) were plated at 5,000 cells per well of a 96-well plate 72hrs before transduction. Crude AAV preps were then diluted 1:1 in DMEM (4.5g/L glucose, Penn/Strep, 20% FBS) for C2C12 cells or Skeletal Muscle Cell Growth Medium for HSkMC cells. Cells were transduced with varying capsid serotypes containing CB7.mScarlet for 24 hours at 37°C. Live cells were stained with Hoechst dye for 10min and imaging was conducted on the Perkin Elmer Opera Phenix using the DAPI (for Hoechst imaging) and mCherry (for mScarlet imaging) filter sets.

## Supporting information

PEI Optimization

Triple Plasmid Transfection

## ACKNOWLEDGMENTS

We thank Robert Tjian and Xavier Darzacq for their support and use of laboratory equipment. We thank Mark Kay for his gift of the KP1 and LK03 rep/cap plasmids, and Luk Vandenberghe for AAV4 rep/cap plasmid. Funding was provided by the Howard Hughes Medical Institute (34430, R. T.) and the California Institute for Regenerative Medicine Training Program EDUC4-12790.

## References

1 Ling, C. et al. The Adeno-Associated Virus Genome Packaging Puzzle. J Mol Genet Med. 9 (3), (2015).

2 Maurer, A. C. & Weitzman, M. D. Adeno-Associated Virus Genome Interactions Important for Vector Production and Transduction. Hum Gene Ther. 31 (9-10), 499–511, (2020).

3 Srivastava, A. Replication of the adeno-associated virus DNA termini in vitro. Intervirology. 27 (3), 138–147, (1987).

4 Wang, X. S., Ponnazhagan, S. & Srivastava, A. Rescue and replication of adeno-associated virus type 2 as well as vector DNA sequences from recombinant plasmids containing deletions in the viral inverted terminal repeats: selective encapsidation of viral genomes in progeny virions. J Virol. 70 (3), 1668–1677, (1996).

5 Earley, L. F. et al. Adeno-Associated Virus Serotype-Specific Inverted Terminal Repeat Sequence Role in Vector Transgene Expression. Hum Gene Ther. 31 (3-4), 151–162, (2020).

6 Yang, J. et al. Concatamerization of adeno-associated virus circular genomes occurs through intermolecular recombination. J Virol. 73 (11), 9468–9477, (1999).

7 Au, H. K. E., Isalan, M. & Mielcarek, M. Gene Therapy Advances: A Meta-Analysis of AAV Usage in Clinical Settings. Front Med (Lausanne). 8 809118, (2021).

8 Zincarelli, C., Soltys, S., Rengo, G. & Rabinowitz, J. E. Analysis of AAV serotypes 1-9 mediated gene expression and tropism in mice after systemic injection. Mol Ther. 16 (6), 1073–1080, (2008).

9 Ellis, B. L. et al. A survey of ex vivo/in vitro transduction efficiency of mammalian primary cells and cell lines with Nine natural adeno-associated virus (AAV1-9) and one engineered adeno-associated virus serotype. Virol J. 10 74, (2013).

10 Atchison, R. W., Casto, B. C. & Hammon, W. M. Adenovirus-Associated Defective Virus Particles. Science. 149 (3685), 754–756, (1965).

11 Meier, A. F., Fraefel, C. & Seyffert, M. The Interplay between Adeno-Associated Virus and its Helper Viruses. Viruses. 12 (6), (2020).

12 McCarty, D. M., Young, S. M., Jr. & Samulski, R. J. Integration of adeno-associated virus (AAV) and recombinant AAV vectors. Annu Rev Genet. 38 819–845, (2004).

13 Yang, Y. et al. A dual AAV system enables the Cas9-mediated correction of a metabolic liver disease in newborn mice. Nat Biotechnol. 34 (3), 334–338, (2016).

14 Bi, X. & Liu, L. F. DNA rearrangement mediated by inverted repeats. Proc Natl Acad Sci U S A. 93 (2), 819–823, (1996).

15 Samulski, R. J., Berns, K. I., Tan, M. & Muzyczka, N. Cloning of adeno-associated virus into pBR322: rescue of intact virus from the recombinant plasmid in human cells. Proc Natl Acad Sci U S A. 79 (6), 2077–2081, (1982).

16 Vandenberghe, L. H. et al. Efficient serotype-dependent release of functional vector into the culture medium during adeno-associated virus manufacturing. Hum Gene Ther. 21 (10), 1251–1257, (2010).

17 Sommer, J. M. et al. Quantification of adeno-associated virus particles and empty capsids by optical density measurement. Mol Ther. 7 (1), 122–128, (2003).

18 Zhu, J., Huang, X. & Yang, Y. The TLR9-MyD88 pathway is critical for adaptive immune responses to adeno-associated virus gene therapy vectors in mice. J Clin Invest. 119 (8), 2388–2398, (2009).

19 Wagner, H. & Bauer, S. All is not Toll: new pathways in DNA recognition. J Exp Med. 203 (2), 265–268, (2006).

20 Sanmiguel, J., Gao, G. & Vandenberghe, L. H. Quantitative and Digital Droplet-Based AAV Genome Titration. Methods Mol Biol. 1950 51–83, (2019).

21 Grimm, D. et al. Titration of AAV-2 particles via a novel capsid ELISA: packaging of genomes can limit production of recombinant AAV-2. Gene Ther. 6 (7), 1322–1330, (1999).

22 Summerford, C. & Samulski, R. J. Membrane-associated heparan sulfate proteoglycan is a receptor for adeno-associated virus type 2 virions. J Virol. 72 (2), 1438–1445, (1998).

23 Bell, C. L. et al. The AAV9 receptor and its modification to improve in vivo lung gene transfer in mice. J Clin Invest. 121 (6), 2427–2435, (2011).

24 McCown, T. J., Xiao, X., Li, J., Breese, G. R. & Samulski, R. J. Differential and persistent expression patterns of CNS gene transfer by an adeno-associated virus (AAV) vector. Brain Res. 713 (1-2), 99-107, (1996).

25 Weitzman, M. D., Kyostio, S. R., Kotin, R. M. & Owens, R. A. Adeno-associated virus (AAV) Rep proteins mediate complex formation between AAV DNA and its integration site in human DNA. Proc Natl Acad Sci U S A. 91 (13), 5808–5812, (1994).

26 Miller, D. G., Petek, L. M. & Russell, D. W. Adeno-associated virus vectors integrate at chromosome breakage sites. Nat Genet. 36 (7), 767–773, (2004).

27 Hanlon, K. S. et al. High levels of AAV vector integration into CRISPR-induced DNA breaks. Nat Commun. 10 (1), 4439, (2019).

28 Porteus, M. H., Cathomen, T., Weitzman, M. D. & Baltimore, D. Efficient gene targeting mediated by adeno-associated virus and DNA double-strand breaks. Mol Cell Biol. 23 (10), 3558–3565, (2003).

